# Endocannabinoid ligands (CBD, Δ9THC, and Terpenes) inhibit excitability of mouse dorsal root ganglion neurons and exhibit synergistic inhibitory effects

**DOI:** 10.64898/2026.07.17.739255

**Authors:** Hedaythul Choudhury, Mazin Nicola, Barnaby W. Greenland, Daniel Guest, John Spencer, Andrew Dilley

## Abstract

The need for improved treatments for chronic pain has driven increased interest in cannabis-based therapeutics. Peripheral dorsal root ganglion (DRG) neurons, including nociceptors, express cannabinoid receptors (CB1 and CB2), suggesting that modulation of DRG excitability may provide an effective strategy for peripheral analgesia. Here, we investigated the effects of cannabidiol (CBD), Δ^9^-tetrahydrocannabinol (THC), terpene mixtures as well as cannabis plant extracts on neuronal excitability in small-diameter mouse DRG neurons using whole-cell current-clamp electrophysiology and assessed potential synergistic interactions. Both CBD and THC produced a concentration- and time-dependent inhibition of rheobase-evoked action potential firing, which were reversible in the presence of bovine serum albumin (BSA), both with similar estimated IC_50_ values of 5 μM (. Terpene mixtures, as well as individual terpenes (linalool, β-pinene, and myrcene), similarly reduced neuronal firing. Co-application of CBD with THC or terpenes enhanced inhibition, consistent with synergistic interactions and the known “entourage effect.” Application of WIN55,212-2 (WIN), a non-selective cannabinoid receptor agonist, in the presence of CBD also accelerated the time-dependent inhibition of neuronal firing. The inhibition of firing by the CB2-selective inverse agonist JTE-907 indicated the presence of CB2 receptors on DRG neurons. Plant extracts from the *Cannabis sativa* leaves also reversibly inhibited neuronal firing. CBD and a terpenes mixture produced modest effects on hERG channels, whereas plants extracts had negligible effects. Collectively, these findings demonstrate that phytocannabinoids and terpenes suppress peripheral sensory neuron excitability via receptor-dependent and indirect mechanisms, supporting their potential as non-opioid analgesics. Their synergistic interactions suggest that multi-component formulations may enhance analgesic effects.

## 1.0 INTRODUCTION

Chronic pain is a major healthcare challenge that significantly impairs quality of life and imposes a substantial socioeconomic burden worldwide. Current treatments, such as anticonvulsants and antidepressants, are often ineffective, and analgesics such as opioids are not recommended for long-term treatment of chronic pain due to risks associated with tolerance and physical dependency[44; 45].

Cannabis-based products have received considerable interest for the management of pain (reviewed in [16]), mostly due to their analgesic properties. To date, more than 500 compounds have been identified from the *Cannabis sativa* plant, including more than 100 phytocannabinoids, as well as terpenes, flavonoids, and other bioactive constituents. Of these, Δ9-tetrahydrocannabinol (THC) and cannabidiol (CBD) are the most extensively studied and pharmacologically active constituents. THC, the primary psychoactive cannabinoid, functions as a partial agonist at cannabinoid receptors CB1 and CB2 [34]. In contrast, CBD is not psychoactive but exhibits a broad pharmacological profile. I t interacts with multiple targets, such as the CB1 receptor, where it acts as a negative allosteric modulator[25; 33], transient receptor potential (TRP) channels[6; 35; 40], and serotonin receptors (5-HT3, 5-HT_1A_)[3]. These interactions are thought to underlie both its analgesic and anti-inflammatory and analgesic properties.

Components of the endocannabinoid system (ECS), such as CB1, CB2, GPR18, and GPR55 receptors, are expressed throughout the central nervous system [19]. Within the periphery, dorsal root ganglion (DRG) neurons, including nociceptive neurons, are shown to express both CB1 and CB2 receptors[10; 17; 38]. As such, modulation of DRG excitability by cannabinoid ligands may provide an effective mechanism for peripheral analgesia in chronic pain conditions[32; 41].

In addition to cannabinoids, terpenes, a class of compounds from the *Cannabis sativa* plant, have recently emerged as bioactive agents with potential analgesic benefits. Common terpenes, such as linalool, β-pinene, and myrcene, have been shown to have analgesic properties[2; 28], and are considered to act on ion channels, such as TRP channels [22]. Importantly, terpenes have been found to modulate cannabinoid effects through the “entourage effect” [7; 8; 14; 26], a synergistic interaction between compounds that enhances therapeutic effects. While the analgesic potential of terpenes is increasingly recognised, electrophysiological studies of their effects on sensory neuron excitability remain limited [5; 21; 27; 39].

Despite substantial interest in the use of cannabis-based products for the treatment of pain, there are currently no FDA-approved drugs for its use in its treatment. However, the FDA has approved the use of CBD formulations for other medical indications (Table 1).

**Table 1:** FDA-approved cannabidiol-based products for different therapeutic indications.

| Drug (Brand name) | Active ingredient | Primary indication(s) |
| --- | --- | --- |
| **Epidiolex** | Cannabidiol<sup>™</sup> (CBD)<br>oral solution | Seizures associated with Lennox-Gastaut syndrome, Dravet syndrome, and tuberous sclerosis complex. |
| **Marinol** | Dronabinol<sup>™</sup><br>(synthetic $\Delta^9$-THC)<br>capsules | Nausea/vomiting from chemotherapy; anorexia with weight loss in AIDS (various historic indications). |
| **Syndros** | Dronabinol<sup>™</sup> oral<br>solution | Same active ingredient (dronabinol) in oral solution form — approved for chemotherapy-induced nausea and other dronabinol indications. |
| **Cesamet** | Nabilone<sup>™</sup> (synthetic<br>cannabinoid<br>analogue) | Chemotherapy-induced nausea and vomiting. |

Multiple studies have examined the effects of CBD and THC in preclinical models of pain (reviewed in [15]). In the present study, we employed whole-cell current-clamp electrophysiology to investigate the effects of cannabinoid-derived compounds, including CBD, THC, terpenes, as well as extracts from *Cannabis sativa* leaves, on rheobase action potential firing in small-diameter mouse DRG neurons (mDRG). We specifically assessed concentration-dependent inhibition, reversibility of effects, and potential synergistic interactions between cannabinoids and terpenes. Together, our findings provide new insight into the mechanisms by which phytocannabinoids modulate sensory neuron excitability and support their potential as non-opioid analgesic agents.

## 2.0 METHODS

### 2.1 Animal Preparation and Tissue Collection

Experiments were performed using 8-week-old C57BL/6J mice housed under standard conditions with access to water and a normal diet. On experimental days, animals were sacrificed in accordance with the Home Office Animal Licensing Schedule 1 Act (Animal Welfare Act 1986). Spinal cords were rapidly removed and placed in ice-cold Hanks’ Balanced Salt Solution (HBSS) without calcium and magnesium (HBSS -/- CaCl_2_ / MgCl_2_, with 0.1% pen strep) dissection buffer. Under a dissecting microscope, each spinal cord was divided into three segments. The dura mater was carefully removed to expose the dorsal root ganglia (DRG) sacs. Individual DRGs were gently dissected from their protective sacs and collected in HBSS buffer.

### 2.2. Reagents

The following reagents were used: CBD (#C-045-1, Cayman Chemical, Sigma; #90080), Δ^9^THC (# T2386, Sigma), WIN55212-2 (#10009023, Cayman Chemical), TEA (#471283, Sigma), Lidocaine (#L7757, Sigma), JTE907 (#10009875, Cayman Chemical), TTX (Abcam), Capsaicin (#HY-10448, Insight Biotechnology), Capsazepine (#HY-15640 Insight Biotechnology), Linalool (#61706 Merck), Myrcene (#6464-3 Merk), β-pinene (#88335 Phytolab), and Terpene Mix-A (# CRM40755, Sigma; Chemical composition (2000 µg/ml of each component in methanol): Camphene, alpha-Pinene, 3-Carene, alpha-Terpinene, (R)-(+)-Limonene, gamma-Terpinene, L-(-)-Fenchone, Fenchol, (1R)-(+)-Camphor, I soborneol, Menthol, Citronellol, (+)-Pulegone, Geranyl acetate, alpha-Cedrene, alpha-Humulene, Nerolidol, (+)-Cedrol, (-)-alpha Bisoprolol).

### 2.3 Extraction of cannabinoid-based plant products

Compounds were extracted from Cannabis sativa (hemp) leaves to generate three full-spectrum extracts: FSE THC80M (Code #002442), FSE CBD-THC (Code #- 002709), and FSE CBD GMP (Code #001778). Plant material was supplied by Clever Leaves (Portugal) under a UK Home Office licence agreement. Extraction and preparation procedures were performed in-house using proprietary methods (IP: GB2610556). Following purification, stock solutions of each extract were prepared in ethanol. The chemical composition of each extract was characterised by high-performance liquid chromatography (HPLC) and mass spectrometry (Table 2, & Supplementary Fig S4).

**Table 2.** Composition of cannabinoid-based plant extracts.

|  |  | Calculated Concentration (ppm) (from Extraction) |  |  |  |  |  |  |  | Ratio |  |
| --- | --- | --- | --- | --- | --- | --- | --- | --- | --- | --- | --- |
| Sample ID | Sample code | CBD | THC | CBDA | CBGA | CBC | CBDV | CBLA | Total (ppm) | THC:CBD | CBD:THC |
| CBD FSE GMP | Extract A | 3500 | 10.37 |  |  |  | 9 |  | 3519 | 0.003:1 | 337.5:1 |
| CBD FSE THC | Extract B | 2445 | 54.9 | 27.25 |  | 89 | 10 | 12 | 2637 | 0.023:1 | 44.5:1 |
| THC 80M | Extract C | 41.34 | 2887 |  | 7 | 41 |  |  | 2948 | 69.8:1 | 0.014:1 |

*Concentrations of individual cannabinoids, expressed as both ppm and mM, were calculated from the mass spectrometry data. The CBD ratio was also calculated for each extract to highlight the marked differences in cannabinoid composition among the three formulations.*

### 2.4 DRG Neuron Isolation and Culture

Following collection, DRG were washed with warm HBSS (#14025092, Thermo Fisher) and subsequently with seeding medium composed of Minimum Essential Medium (MEM) (#912492013, Invitrogen), 0.1% penicillin/streptomycin (#P4333, Sigma), 10% heat-inactivated foetal bovine serum (#16140071, Thermo Fisher), 1% sodium pyruvate (#11360070, Thermo Fisher), and 2.5% HEPES 1M (#15630056, Thermo Fisher). Between washes, tissues were centrifuged at 1200 rpm for 7 minutes. DRGs were enzymatically digested (Collagenase IV and Dispase-II), incubated in digestion medium containing Pluri-Stem Dispase-II (#SCM133M, Speciality Media, Germany) and 0.125% collagenase IV (#C9407, Sigma) for 30 minutes at 37 °C in a water bath. Following digestion, tissues were gently triturated using a Gilson pipette to create a cloudy single-cell suspension.

The cell suspension was washed three times and centrifuged at 1200 rpm for 7 minutes between each wash. Finally, the supernatant was removed, and the cell pellet was resuspended in seeding medium. The suspension was filtered through a cell strainer, and the volume was adjusted for seeding. Sterile coverslips were prepared 48 hours before DRG isolation by coating with 0.1 mg/ml Poly-D-lysine (#P4707, Sigma) for 4-6 hours at room temperature, followed by washing, air-drying, and overnight coating with 2 μg/ml laminin (#L2020, Sigma) at 4 °C. On the day of seeding, laminin was removed, and approximately 10,000 cells per well were seeded and incubated for 1 hour at 37 °C with 5% CO_2_. Culture medium containing Neurobasal™-A medium (#100888-022, Thermo Fisher), 10 ng/ml recombinant mouse NGF (#13257-019, Thermo Fisher), 1× B27 supplement (#17504044, Thermo Fisher), 1% penicillin/streptomycin (#P4333, Sigma), and 2 mM L-glutamine (#A2916801, Thermo Fisher) was then added to each well. Medium was changed (50:50 replacement) after the first 24 hours and every 48 hours thereafter, supplemented with NGF.

### 2.5 Whole-Cell Electrophysiology

Current clamp recordings were performed in whole-cell configuration at room temperature (20 °C) using an Axopatch-700B amplifier (Axon Instruments, CA, USA). DRG neurons were selected for e-phys study based on their size and were generally <50 µM in diameter. Current recordings were low-pass filtered at 10 kHz using a Bessel filter and sampled at 20 kHz with a Digidata 1550B data acquisition system (Axon Instruments, USA). Data acquisition and command voltages were controlled using Clampex 11.2 software (Axon Instruments, USA). Patch pipettes were fabricated from borosilicate glass capillaries (Warner Instruments, UK) using a P-97 Flaming/Brown Micropipette Puller (Sutter Instruments, CA, USA) and heat-polished with an MF900 Microforge (Narishige, Japan) to achieve resistances of 2.5-5 MΩ. Pipettes were filled with intracellular solution containing (in mM): 60 KCl, 20 KF, 60 K-gluconate, 2 Mg-ATP, 10 EGTA, 1.5 MgCl_2_, 1 CaCl_2_, and 10 HEPES, with pH adjusted to 7.32 using KOH and osmolarity adjusted to 320 mOsm with sucrose. Pipette solutions were prepared in batches, aliquoted, and stored at −20°C until use. All solutions were equilibrated to room temperature before the experiments.

DRG neurons were visualized using an inverted microscope (Nikon TSE 300, Japan) with 40× objectives and continuously superfused with extracellular bath solution (ECS) containing (in mM): 140 NaCl, 5 KCl, 2 MgCl_2_, 2 CaCl_2_, 10 HEPES, and 10 D-glucose, with pH adjusted to 7.35 using NaOH and osmolarity adjusted to 300 mOsm. The recording chamber was grounded with an Ag/AgCl pellet, and junction potentials between intracellular and extracellular solutions were zeroed with the filled pipette in the bath solution. Neurons were selected for recording based on resting membrane potential values between −45 and −60 mV and action potential overshoot exceeding +20 mV. After achieving whole-cell configuration, cells were held in current clamp mode, and the resting membrane potential was recorded. Rheobase current, defined as the minimum current required to elicit action potentials, was determined using a series of current step protocols. Once steady baseline firing was established, vehicle and test compounds were applied sequentially using a fast perfusion system (RSC200 Rapid Perfusion system with EVH-9 rapid valve system, Biologic Science Instruments, France). A reference compound was applied at the end of each experiment for comparison and normalisation purposes. Stock solutions were prepared in 100% DMSO (Sigma) or appropriate vehicle according to the manufacturer’s instructions. Working concentrations were prepared by diluting stock solutions in ECS to achieve a final DMSO concentration of 0.3%.

Owing to the limited aqueous solubility and physicochemical properties of CBD and THC, full log dose-response curves were not feasible. Instead, 2- and 3-point concentration–response experiments were conducted using 2-fold concentration increments.

Drugs were applied sequentially in the following order: ECS, vehicle control (ECS containing BSA), and the test compound. At the end of each recording, a reference compound was applied to confirm assay performance, and responses were normalised to the reference response. All compounds were superfused continuously until a steady-state (equilibrium) response was achieved. Prior to each drug application, the corresponding vehicle control (buffer without the test compound) was applied to establish a baseline response. Drug-induced effects were subsequently normalised to the corresponding control values.

### 2.6 Comprehensive in vitro pro-arrhythmia ion channel screening

Comprehensive in vitro pro-arrhythmia ion-channel screening was conducted to evaluate the cardiac safety profile of CBD ligands and assess their potential off-target effects on cardiac ion channels. Qpatch electrophysiology assays were carried out using recombinantly expressed hNav1.7 co-expressed with β1 and β2 subunits (hNav1.7 β; HEK293) and human Ether-à-go-go-Related Gene (hERG; CHO) cell lines, with TTX and quinidine used as reference compounds for Nav and hERG channels, respectively.

### 2.7 Data Analysis

Initially, drug effects were quantified by measuring peak action potential amplitude (mV), resting membrane potential (mV) and normalising to the vehicle condition immediately preceding drug application (post /pre drug ratio; control = 1). The action potential peak ratio was calculated as the peak amplitude at the end of the final sweep divided by the average peak amplitude of 3–5 action potentials immediately before drug application. Due to variability associated with this approach, the analysis was revised. In subsequent experiments, peak action potential amplitudes were analysed as paired measurements within individual cells, comparing pre-drug and post-drug values directly.

Data analysis was performed using Clampfit 11.2 software (Axon Instruments, USA), with key parameters including resting membrane potential (RMP, mV) and peak action potential amplitude (mV). Data were normalised as ratios (control/response) and presented as mean ± SEM (n). Prism 10.6 software was used to generate graphs. Statistical analyses were performed using Prism 10 software with one-way ANOVA for multiple comparisons and unpaired Student t-tests for two-group comparisons, with statistical significance set at p < 0.05.

## 3.0 RESULTS

### 3.1 Pharmacological Properties

Current clamp recordings were used to examine stepwise current-dependent action potential discharge. No action potentials were elicited at negative current steps, while firing was observed only when step currents exceeded +50 pA, without changes in resting membrane potential (Fig.1A, B; Table 2). With 100 pA current injection, tonic firing and phasic firing patterns were observed (Fig. 1C-E). These cells exhibited action potential peak amplitudes between +20 and +40 mV. For bioactive compound testing, both tonic and phasic firing neurons were selected with an RMP less than −40 mV and a peak action potential amplitude greater than +20 mV. These cells were sufficiently stable and enabled longer electrophysiological recording times.

Lidocaine and TTX were used to characterise action potential properties (Fig. 1F-I). Baseline firing was rapidly and reversibly inhibited by both 300 μM lidocaine and 100 nM TTX without affecting the RMP (p<0.001; Fig. 1. F-I). Whole-cell current recordings also demonstrated blockade of outward currents by 10 mM TEA (Supplementary Fig. S1).

**Table 2:** Summary of action potential properties.

| | **RMP (mV)**<br>Mean ± SEM (n) | **AP Threshold (mV)**<br>Mean ± SEM (n) | **Peak Amplitude (mV)**<br>Mean ± SEM (n) |
| --- | --- | --- | --- |
| I-V protocol<br>(200 pA) | -53.72 ± 1.24 (58) | -32.94 ± 1.16 (52) | 39.85 ± 2.52 (55) |
| Single Step protocol<br>(100 pA) | -56.81 ± 1.33 (56) | -33.82 ± 1.32 (55) | 27.28 ± 2.05 (49) |

**Fig. 1.**
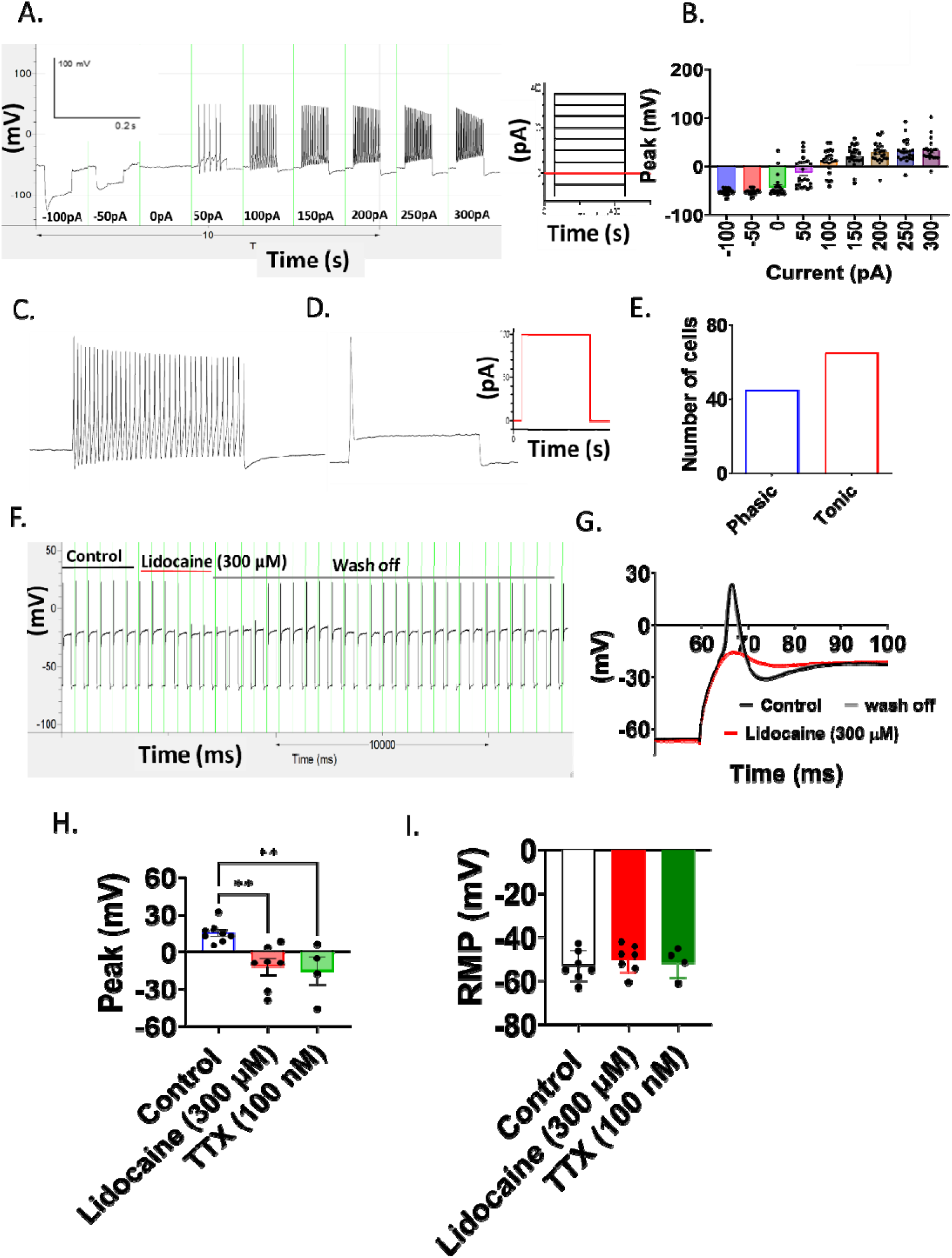
Pharmacological properties of small diameter DRG neurons. (A) Example trace showing action potential discharge from the stepwise current. No action potential discharge was recorded when the current was at a negative potential, and firing was observed when the step current was greater than >+50 pA. (B) Peak action potential (AP) with increasing IV current injection. (C-D) Examples of(C) tonic (burst) and (D) phasic (single) action potential activation at 100 pA current. (E) Number of cells with tonic or phasic action potential discharge. (F) An example recording showing inhibition of baseline firing by 300 μM lidocaine. (G) Overlaid traces showing action potentials in control and in the presence of 300 µM lidocaine. (H-I) Effect of 300 µM lidocaine (n=7) and 100 nM TTX (n=4) on (H) peak action potential (AP) height and (I) resting membrane potential (RMP). Data presented as mean ± SEM (Individual data points overlaid). **p<0. 01.

### 3.2 Effect of CBD and THC on small-diameter DRG neurons

Both CBD and THC caused a concentration-dependent block of neuronal firing without affecting RMP. CBD showed a lower threshold for inhibition, with effects becoming apparent at concentrations of ≥2 µM (Fig. 2A-D), with maximum block observed at these concentrations. The maximum block occurred at >5 minutes following application of the highest dose. Some variability was noted in the effect of CBD on neuronal firing. In contrast to CBD, THC required higher concentrations to produce inhibitory effects (≥ 10 µM), with maximum block observed at 30 μM (Fig. 2E-H). The effects of THC at 10 μM were variable. The maximum block occurred after 2-4 minutes following application of the highest dose. Both CBD and THC caused a progressive time-dependent reduction in action potential amplitude during application. Inhibition of neuronal firing by CBD and THC was not reversible when the compounds were prepared in an extracellular buffer containing either 0.3% DMSO or 0.1% ethanol (Supplementary Fig. 2). The IC_50_ values for both CBD and THC were comparable (∼5 μM) and did not affect the RMP (Fig. 2F, H).

**Fig. 2.**
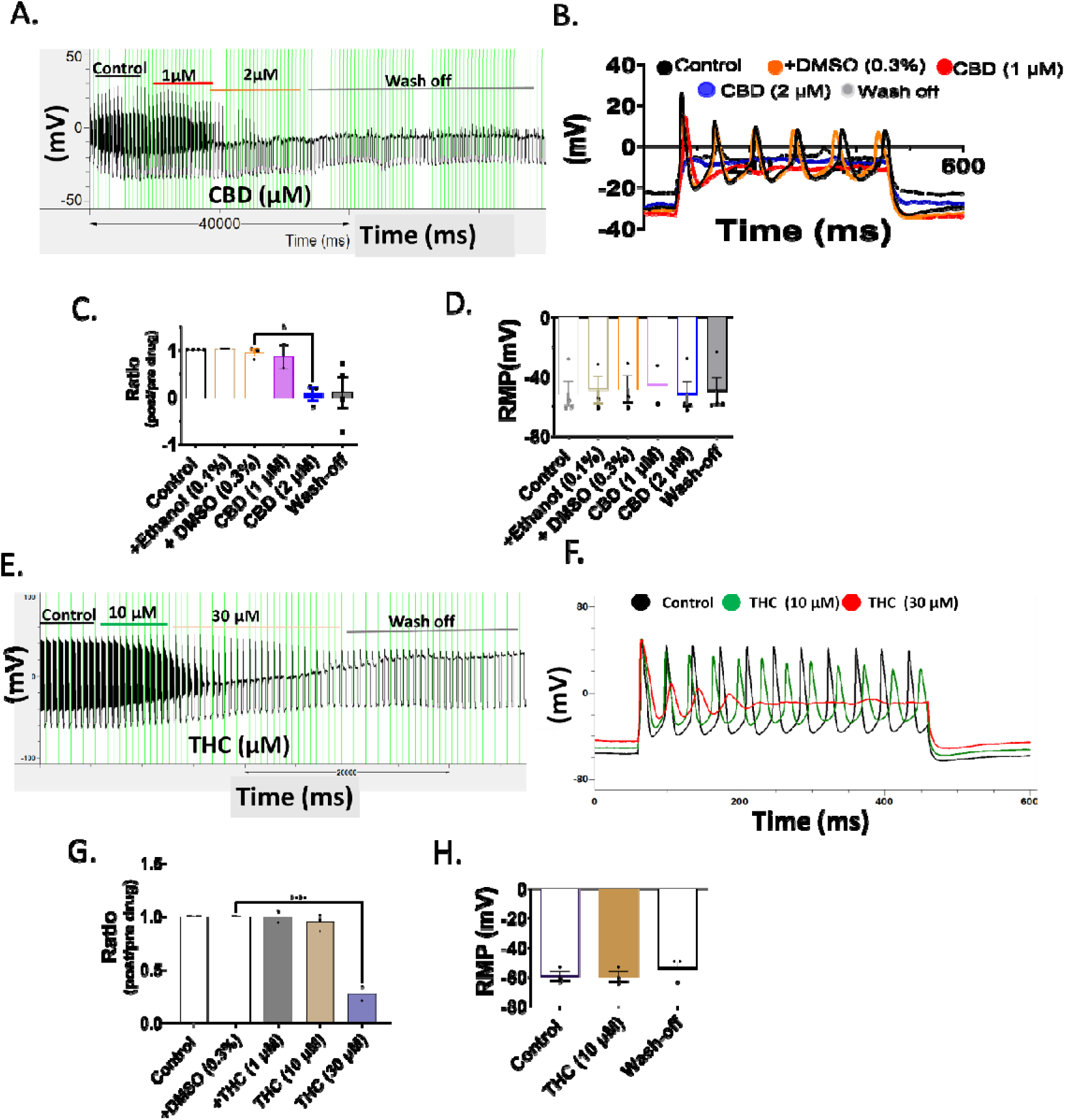
Effects of CBD and THC on neuronal firing. (A) Example trace showing the effect of 1 and 2 µM CBD on neuronal firing. (B) Overlaying sweeps from the same recording. In this example, 2 µM CBD completely blocked neuronal firing. (C) Example trace showing the effects of 10 and 30 µM THC on neuronal firing. (D) Overlaying trace from the same recording. Note the time-dependent progressive reduction in action potential amplitude during the application of CBD and THC. For both CBD and THC, the inhibition of firing was irreversible during the wash-off with 0. 3% DMSO. (E-H) Summary graphs showing the effects of (E, F) CBD and (G, H) THC on (E, G) action potential amplitude (expressed as a ratio) and (F, G) resting membrane potential (RMP; n = 3-4). During washes, no changes in RMP were observed. Data presented as mean ± SEM (Individual data points overlaid). * p<0. 05, **p<0. 01, ****p<0. 0001.

The addition of 0.5% BSA to the extracellular buffer solution led to a concentration-dependent reversible inhibition of neuronal firing by CBD (Fig. 3A, B, E) and THC (Fig. 3C, D, G), with no change in RMP (Fig. 3F, H). Control experiments confirmed that BSA does not affect the neuronal firing or RMP (Fig. 3I, J). Note that, due to the reversibility of CBD and THC in the presence of BSA, all subsequent experiments were conducted using a combined BSA/DMSO vehicle.

**Fig. 3.**
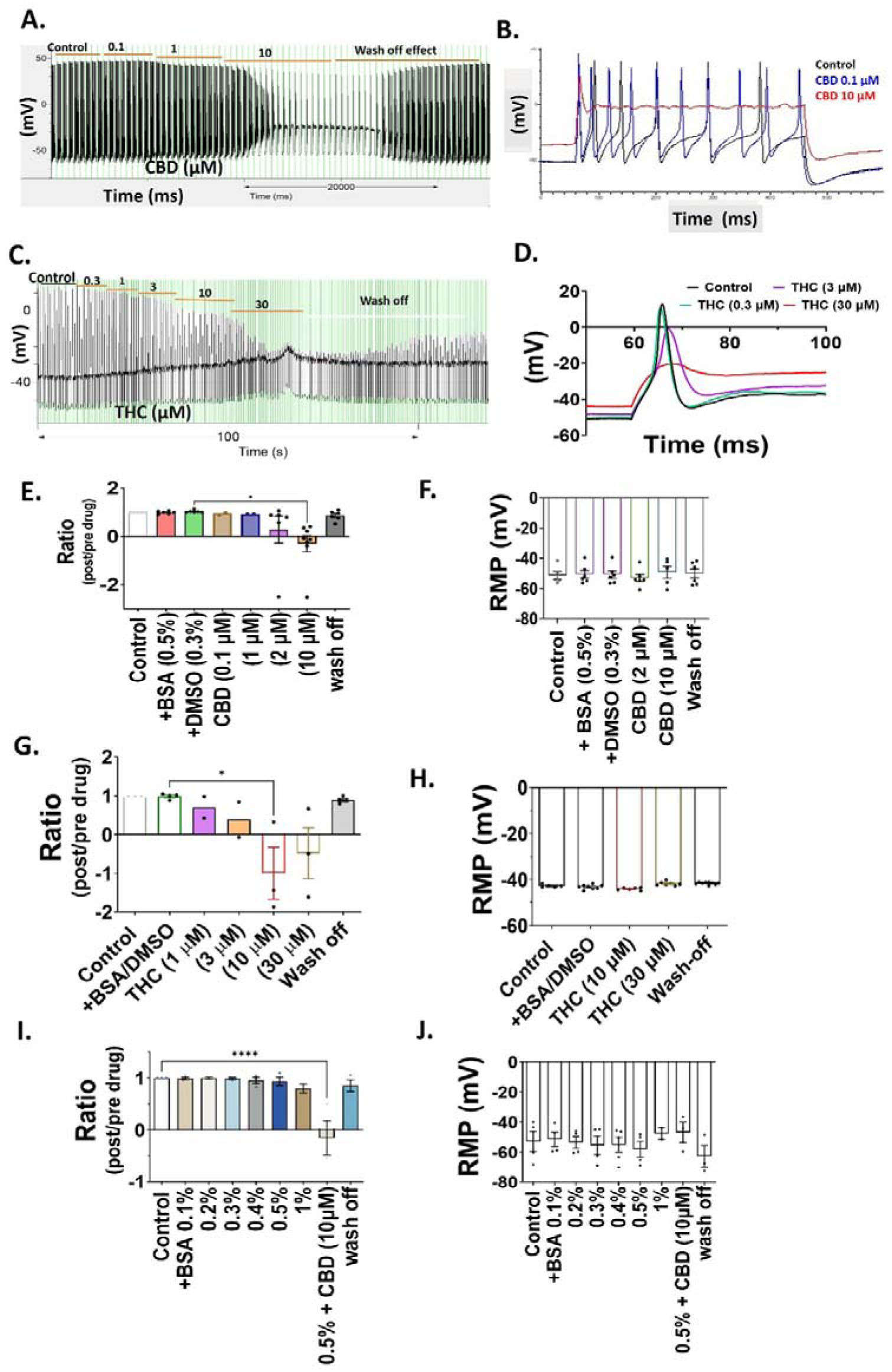
Concentration responses of CBD and THC on neuronal firing in the presence of 0. 5% BSA. (A) Example trace showing the reversible effects of 10 µM CBD in 0. 5% BSA on neuronal firing. (B) Overlaying sweeps from the same recording. (C) Example trace showing the reversible effects of 30 µM THC in 0. 5% BSA on neuronal firing. 0. 3-30 µM THC caused a concentration-dependent block of neuronal firing with a progressive reduction in action potential amplitude. In this example, 30 µM THC was only partially reversible with erratic action potential firing. (D) Overlaying sweeps from the same recording. (E-J) Summary graphs showing the reversible effects of CBD, THC in 0. 5% BSA and BSA alone on (E, G, I) peak action potential amplitude (expressed as a ratio) and (F, H, J) resting membrane potential (RMP; n = 3-6). Note the small shift in RMP in the presence of 1% BSA. Data presented as mean ± SEM (Individual data points overlaid) * p < 0. 05, **** p < 0. 0001.

### 3.3 Synergistic effects of CBD and THC on neuronal firing

To assess the “entourage effect” of CBD and THC, they were assessed together (Fig. 4). Since robust inhibition of neuronal firing was typically observed at CBD concentrations ≥2–5 µM and at THC concentrations ≥5-10 µM, these effective ranges were used to select concentrations for subsequent synergy experiments. Co-application of 5 μM CBD and 10 μM THC produced a reversible block of neuronal firing without altering action potential waveform (Fig. 4A-B). The onset of block was faster (<1 min) than with CBD or THC alone (3–4 min), but slower than with 300 μM lidocaine (<10 s; Fig. 4C–E). In combination, CBD/THC produced a consistent reduction in peak action potential amplitude, whereas CBD or THC alone showed variable effects (Fig. 4F). Resting membrane potential was unchanged under all conditions (Fig. 4G).

**Fig. 4.**
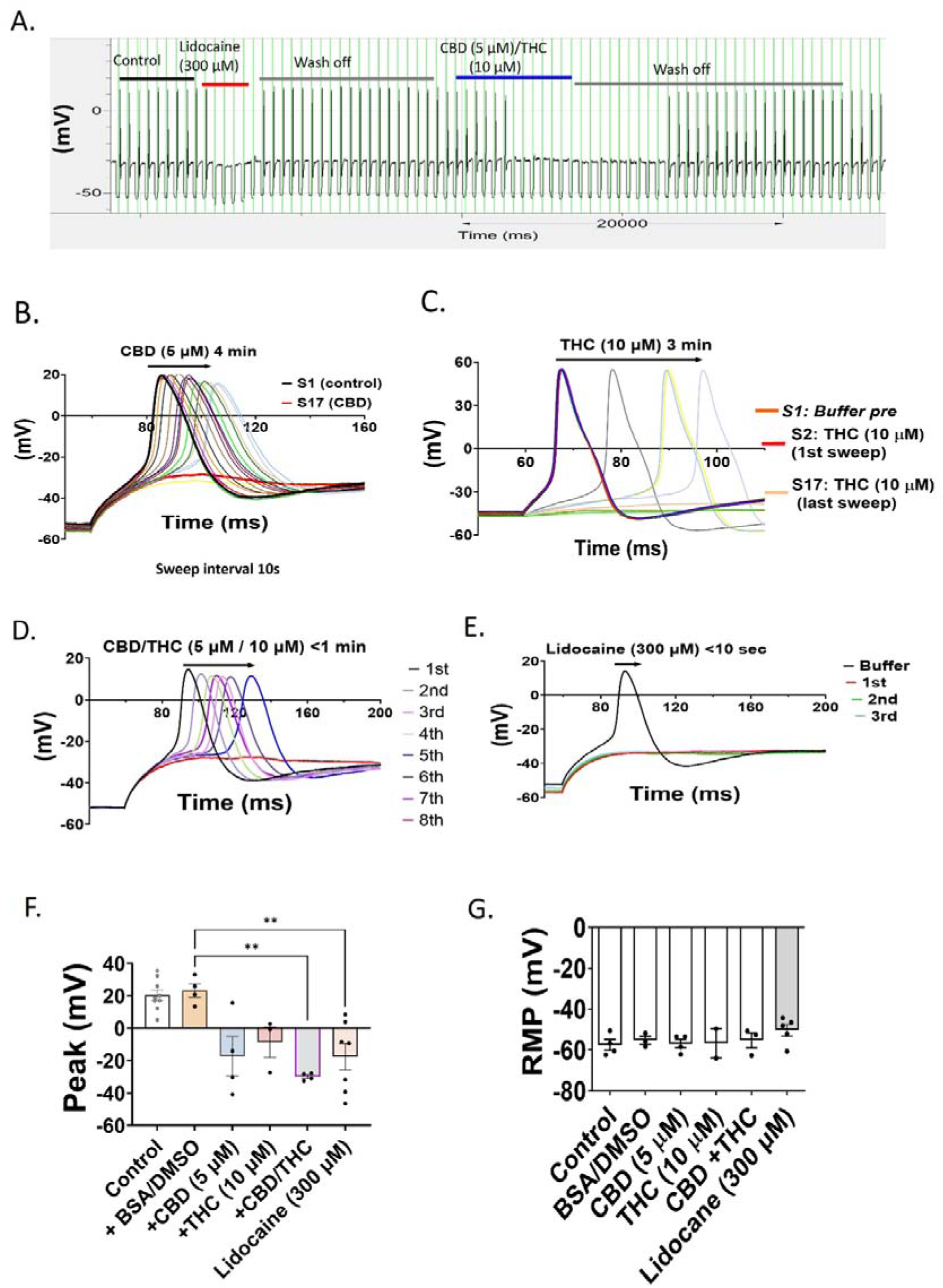
Synergistic effects of CBD and THC on neuronal firing. (A) Example trace showing the effects of 300 µM lidocaine and 1:2 ratio CBD: THC (5 µM CBD + 10 µM THC) on neuronal firing. (B) Overlaid traces of the same recording. (C-F) Kinetic profile of (C) 5 µM CBD, (D) 10 µM THC, (E) 5 µM CBD/10 µM THC and (F) 300 µM lidocaine on the inhibition of neuronal firing. Individual action potential traces are at 10sec intervals. In C and D, 5 µM CBD and 10 µM THC caused a slow (3-4 min) time-dependent complete block of neuronal firing. In D, the complete block of neuronal firing by 5 µM CBD/10 µM THC occurred within 1 min of application, without a progressive reduction in action potential amplitude. In F, 300 µM lidocaine caused an immediate block. (G, H) Summary graphs showing the effects of 5 µM CBD, 10 µM THC, 5 µM CBD/10 µM THC combined and 300 µM lidocaine on (G) peak action potential amplitude and (H) resting membrane potential (RMP). The RMP was not affected by CBD/THC combined. (I-J) Ratiometric analysis showing the effects of increasing [THC] when combined with 2 µM CBD on (I) peak action potential amplitude and (J) RMP (n = 2-5). The optimum effect was with 1:5 CBD: THC ratio. All washes with 0. 3% DMSO/ 0. 5% BSA. Data presented as mean ± SEM (Individual data points overlaid). * p < 0. 05, ** p < 0. 01.

A ratiometric analysis using 2 μM CBD with increasing THC concentrations identified a maximal effect at a 1:5 CBD:THC ratio (Fig. 4I), with no change in resting membrane potential (Fig. 4J). Whole-cell recordings showed that this ratio reduced outward currents by ∼50%, whereas CBD or THC alone did not affect inward or outward currents (Supplementary Fig. 3).

### 3.4 Contribution of Synthetic CB Ligands to the Inhibition of Neuronal Firing

JTE907, a high-affinity CB2 receptor inverse agonist, caused a partially reversible time-dependent reduction in action potential amplitude during application, at 10 μM concentration (Fig. 5A-B), without affecting the RMP (Fig. 5C). The application of 100 nM WIN55,212-2 (WIN), a non-selective cannabinoid receptor agonist, in the presence of 10 μM CBD accelerated the time-dependent inhibition of action potential firing by CBD compared to 10 μM CBD alone (Fig. 5D-F). Preincubation with 100 nM WIN did not affect neuronal activity. The RMP was unaffected by WIN (Fig. 5G).

**Fig. 5.**
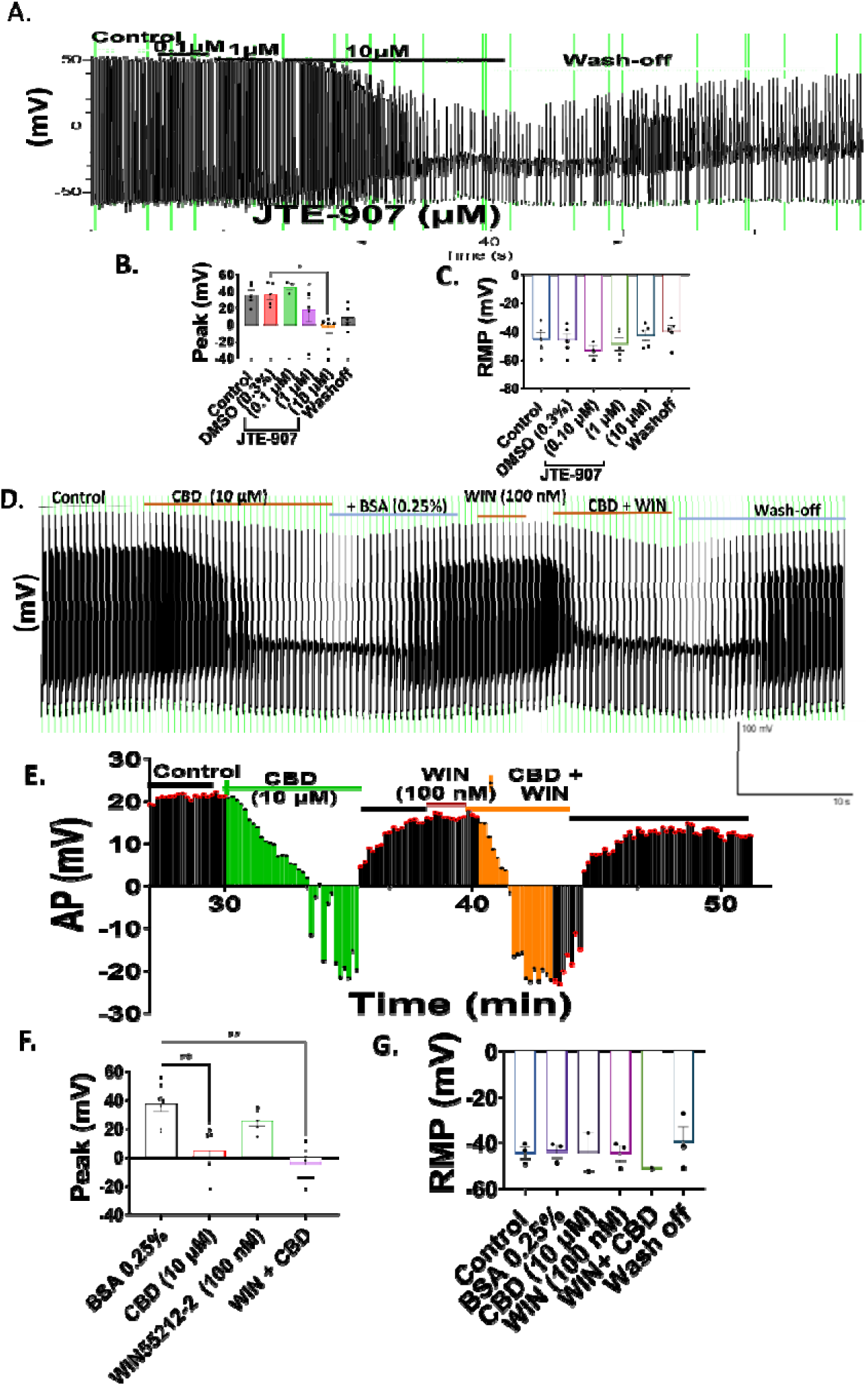
Effects of cannabinoid ligands on neuronal firing. (A-D) Effects of JTE907, an inverse CB2 agonist, on neuronal firing. (A) Example trace showing the effects of 0. 1 µM, 1 µM and 10 µM JTE907. (B) Overlaid traces of the same recording. (C-D) Summary graphs showing the (C) peak action potential amplitude and (D) RMP. (E-I) Effects of 100 nM WIN55,212-2 (WIN), a non-selective cannabinoid receptor agonist on neuronal firing. (E) Example trace showing the reversible effects of 10 µM CBD and 10 µM CBD/100 nM WIN combined. (F) Overlaid traces of the same recording. (G) Kinetics profile comparing 10 µM CBD, 100 nM WIN and 10 µM CBD/100 nM WIN combined. Note the more rapid block by CBD/WIN combined compared to CBD alone. When WIN is applied without CBD, it has no effect. (H-I) Summary graphs showing the (H) peak action potential (AP) amplitude and (I) resting membrane potential (RMP; n = 3-5). All washes with 0. 3% DMSO/ 0. 5% BSA. Data presented as mean ± SEM (Individual data points overlaid). *p<0. 05, **p<0. 01, ***p<0. 001.

### 3.5 Effect of Terpenes on Neuronal Firing

A commercially sourced terpene mixture A containing 19 commonly identified terpenes (Terpenes Mixture-A 5 µg/ml) blocked neuronal firing (Fig. 6A-B). This inhibitory effect was fully reversible (Fig. 6A) and did not affect RMP (Fig. 6C). The vehicle control (0.125% methanol) did not alter baseline action potential firing or the resting membrane potential.

**Fig. 6.**
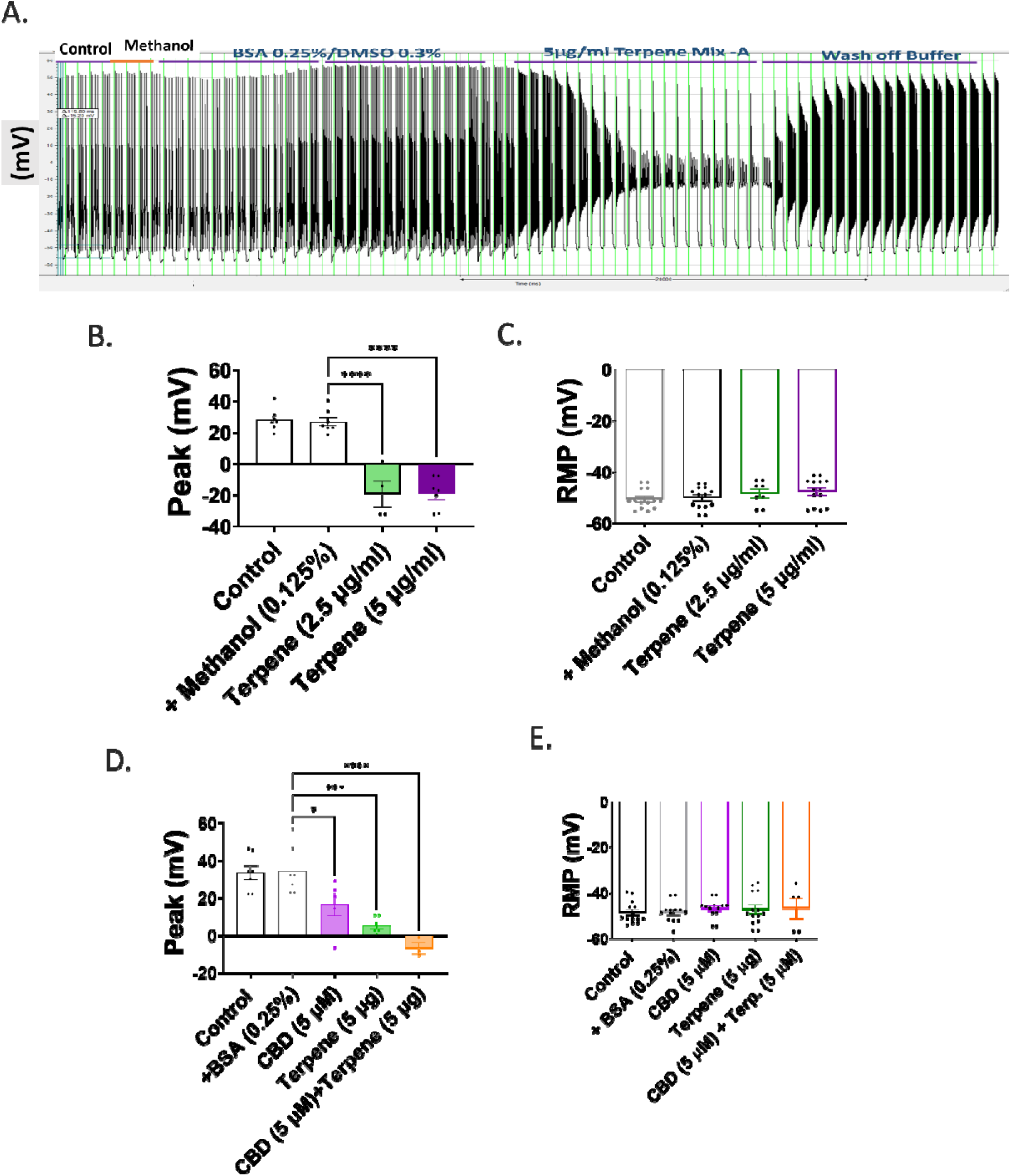
The effect of Terpenes Mixture-A on neuronal firing. (A) Example trace showing block by 5 µg/ml of Terpenes Mixture A. (B) Overlaid traces of the same recording. (C-D) Summary graphs showing the (C) peak action potential amplitude and (D) RMP (n = 4-7). The effects of 0. 125% methanol (vehicle) are also shown. (E and F) Ratiometric analysis showing the synergistic effect between CBD and Terpenes Mixture-A on (E) peak action potential amplitude (AP) and (F) resting membrane potential (RMP; n = 2-7). All washes with 0. 25% DMSO/ 0. 5% BSA. Data presented as mean ± SEM (Individual data points overlaid). *p<0. 05, ***p<0. 001, ****p<0. 0001.

The synergistic effects of Terpenes Mixture-A and CBD were assessed. Terpenes Mixture-A, 5 µg/ml, in the presence of 2 μM CBD produced a concentration-dependent inhibition of neuronal firing (Fig. 6D), without affecting the RMP (Fig. 6E). The effect was more pronounced compared to 5 μg/ml of Terpenes Mixture-A alone (Fig. 6D).

The effects of three common terpenes, linalool, β-pinene and myrcene, on neuronal firing were tested (Fig. 7). Linalool significantly reduced the action potential amplitude (Fig. 7A). Both β-pinene and myrcene also produced a small reduction in amplitude, although the effect was variable between experiments (Fig. 7 C. E). The reduction of neuronal firing was irreversible. None of the terpenes affected the RMP (Fig. 7B, D, F).

**Fig. 7.**
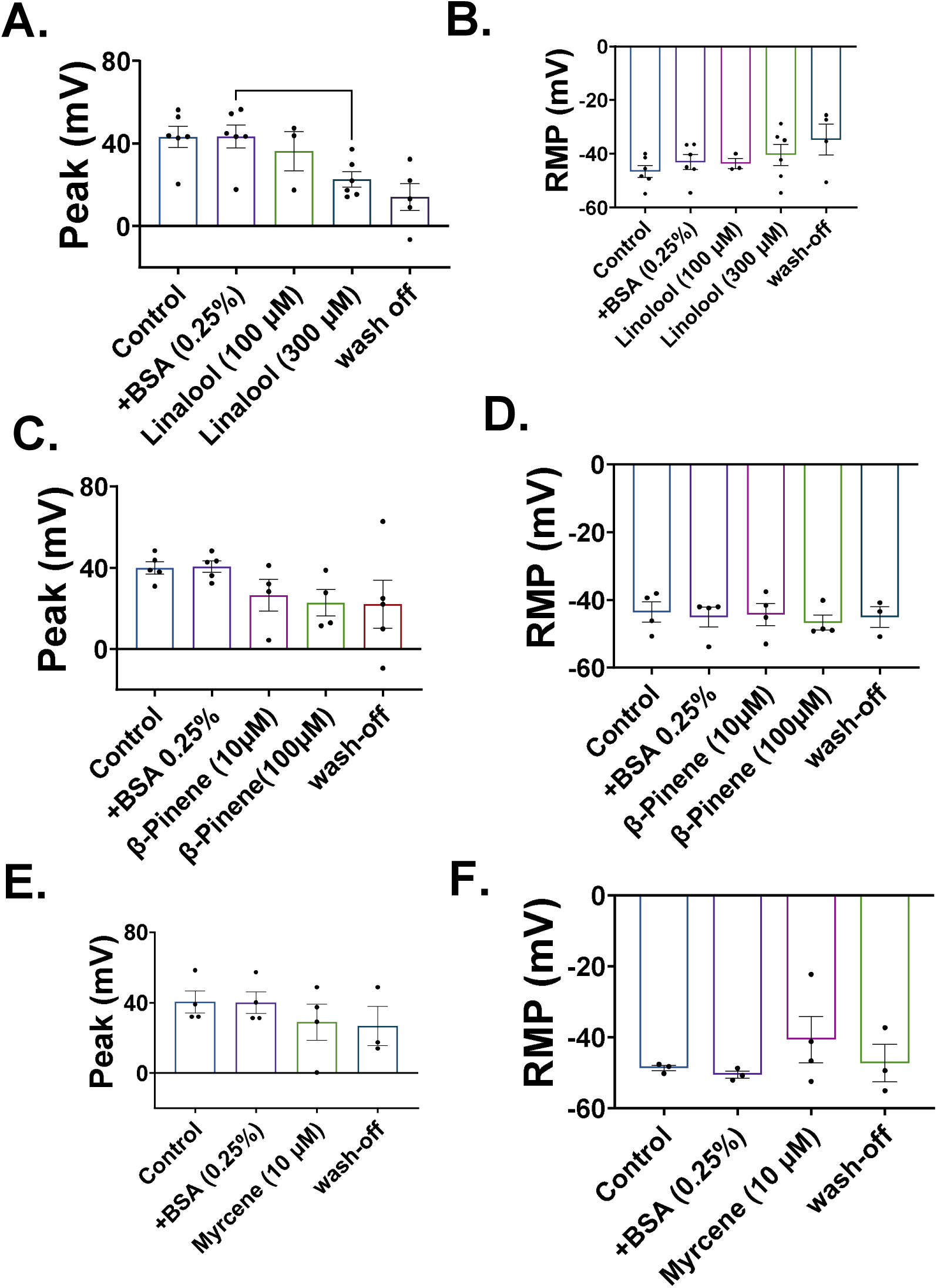
The effects of individual terpenes on neuronal firing. The effects of (A, B) linalool (n=xx), (C, D) β-pinene (n=xx), and (E, F) myrcene (n=) on (A, C, E) peak action potential amplitude and (B, D, F) RMP. For both β-pinene and Linalool, higher concentrations were required to block the current compared to myrcene, which acted at much lower concentrations. The inhibition by individual terpenes failed to reverse following wash-off. Data shown are mean ± SEM (Individual data points are also shown; n = 4). * p < 0. 05.

### 3.6 Effect of natural cannabinoid plant extracts on rheobase current

All three plant extracts were prepared in a combination of methanol/BSA, which inhibited rheobase-evoked neuronal firing, with maximal effects observed at 10 µM (Fig. 8A-F). The composition of the plant extracts is summarised in Table 2. The ratio of CBD:THC in extracts A, B and C was 337.5:1, 44.5:1 and 0.014:1, respectively. Inhibition developed rapidly for all three extracts, reaching a maximum effect within 3–4 min of application, and was partially reversible following washout. None of the extracts significantly altered RMP (Fig. 8G-I). Similar to purified CBD and THC, the extracts produced non-logarithmic concentration-dependent effects (Fig. 8J-K). The IC_50_ for extracts B and C was lower compared to CBD, THC and extract A (Fig. 8L). Comparing ratios (post/pre drug), all the extracts had a similar effect at 10 µM (Fig. 8M).

**Figure 8:**
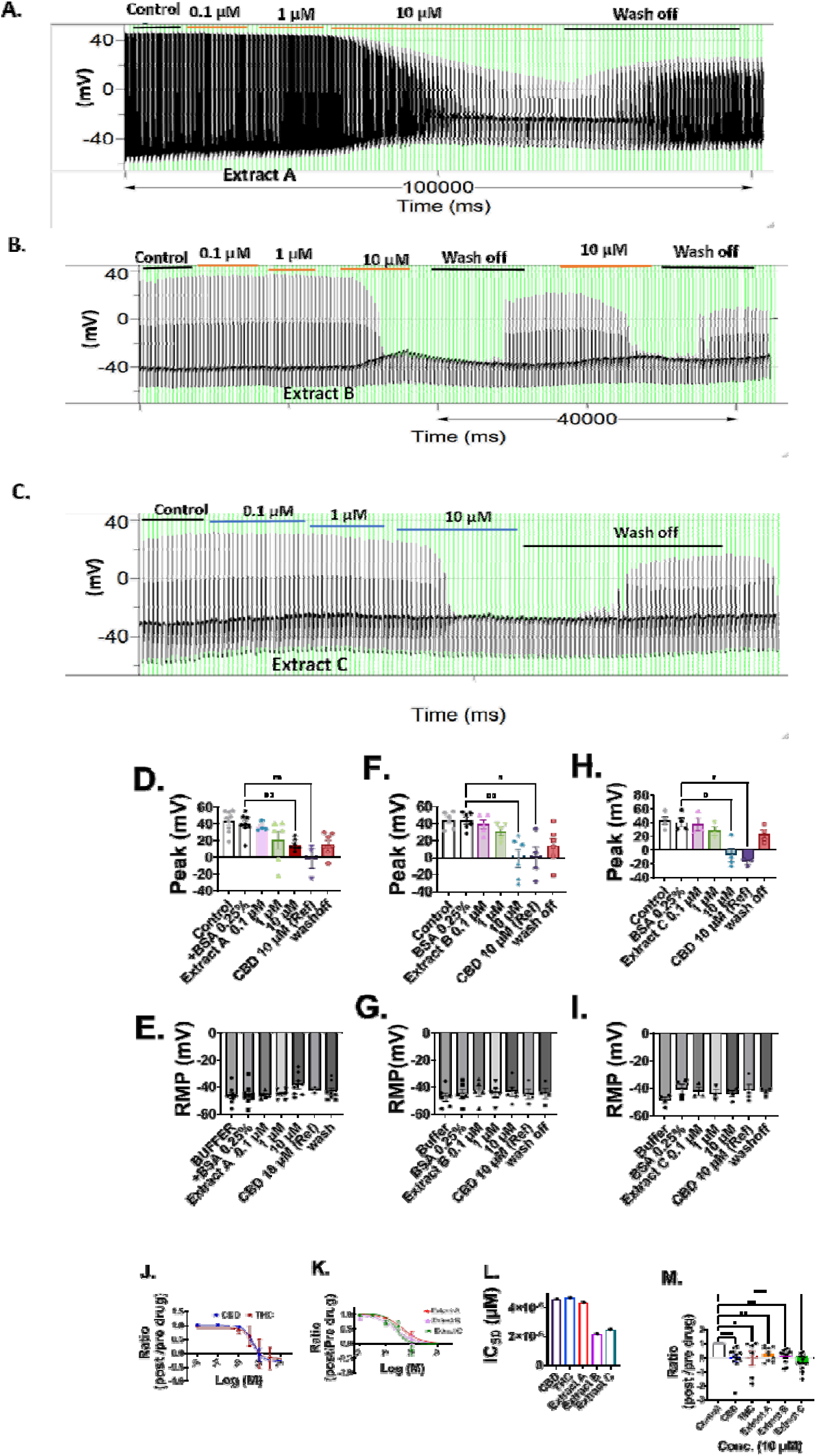
Examples of current clamp recording showing the effect of different plant extracts. (A-C) Example traces showing the effects of 0. 1-10 µM of Extract A, B and C on neuronal firing. (D-I) Summary graphs showing the effects of each extract on (D-F) peak action potential amplitude and (G-I) resting membrane potential (RMP). (J-K) Concentration–response curves for synthetic (J) CBD and THC, and (K) each extract. Curves were fitted using a sigmoidal dose–response model. (L) Summary bar graph showing the IC_50_ values (µM) for the synthetic cannabinoids and plant extracts, calculated from the concentration–response curves. (M) Comparison of the inhibitory effects of the compounds at 10 µM, expressed as a ratio. Data are presented as mean ± SEM. * p <0. 5, ** p < 0. 01, *** p < 0. 001, **** p < 0. 0001.

### 3.7 Effects of cannabinoids and related compounds on Nav1.7β and hERG channels

Using Qpatch electrophysiology assays, the effects of CBD, THC, Terpenes Mixture-A and the plant extracts were examined on Nav1.7β (Fig. 8A-C) and hERG (Fig. 8D-H). There were no significant inhibitory effects by these drugs on Nav1.7β currents either individually or in combination (Fig. 8A-C). 10 µM CBD and the terpene mixture significantly reduced both the step and tail component of the hERG current (Fig. 8G-H). None of the plant extracts significantly affected these channels.

**Fig. 8:**
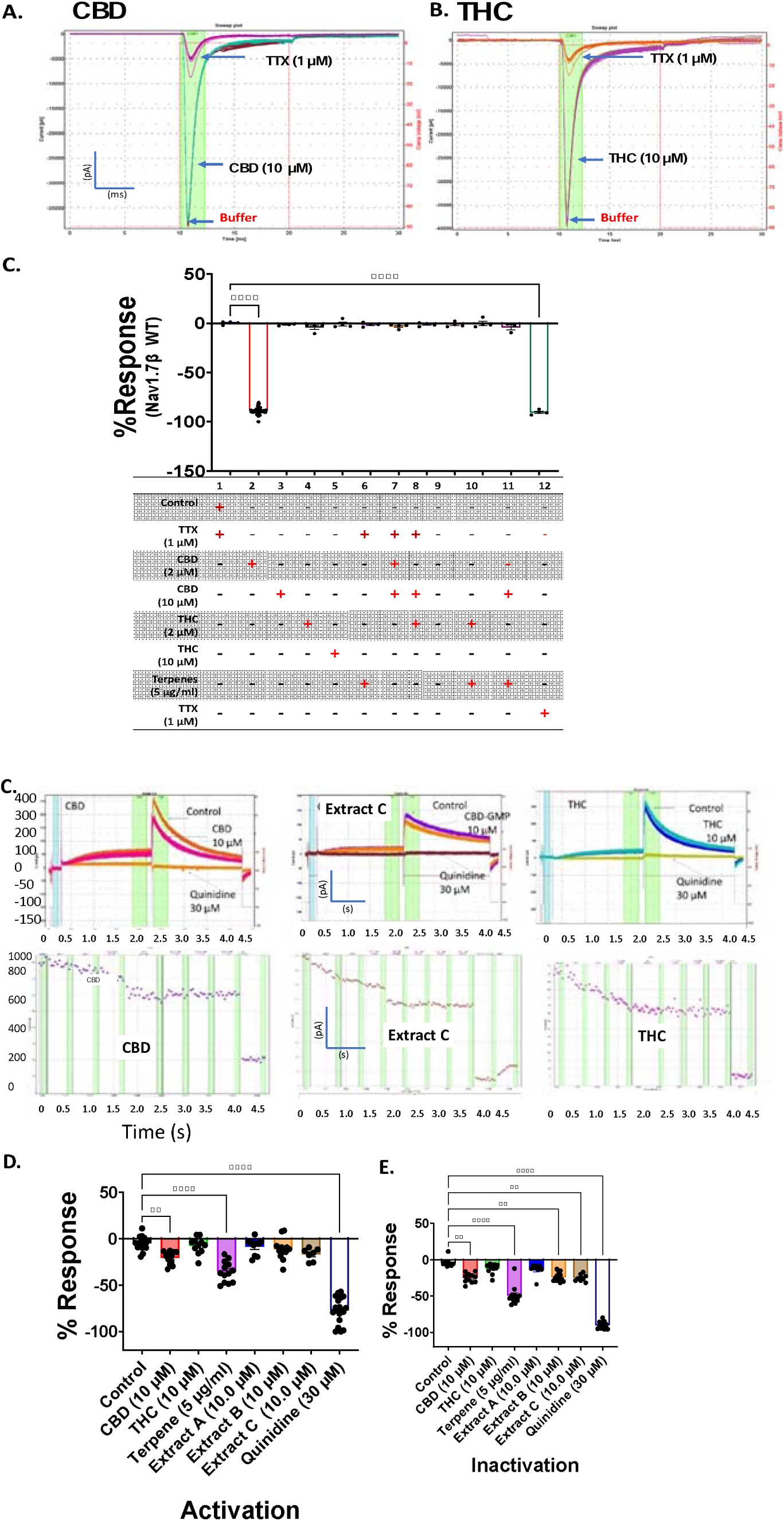
Off-target effect of the cannabinoid ligands on Nav1. 7β and hERG. (A-B) Example traces showing the effect of (A) 10 µM CBD and (B) 10 µM THC, as well as 1 µM TTX on Nav1. 7β (HEK293) channels. (C) Percent response for each ligand (Nav1. 7β; n = 4). (D-F) Example trace showing the effect of (D) 10 µM CBD and (E) 10 µM THC, as well as (F) 10 µM extract C on hERG channels. (G, H) Percent response for each ligand on both the (G) step and (H) tail hERG currents (n= 4). Percent response values represent the percent block relative to the reference compounds (TTX for Nav1. 7β and quinidine for hERG). Data presented as mean ± SEM (Individual data points overlaid). **p<0. 01, ****p<0. 0001.

## 4.0 Discussion

This study demonstrates that CBD, THC, and terpene compounds inhibit neuronal firing in mouse dorsal root ganglion (DRG) neurons with diameters <50 µm. These small-diameter neurons include nociceptive subtypes involved in pain transmission [9; 12]. Using whole-cell current-clamp recordings, all three classes of compounds suppressed rheobase-evoked action potential firing in a concentration- and time-dependent manner. Inhibition was enhanced when cannabinoids and terpenes were co-applied, consistent with a synergistic, termed “entourage effect [7].” These findings indicate that phytocannabinoids and terpenes modulate peripheral nociceptive signalling and may have therapeutic relevance for pain management.

DRG neurons exhibited both tonic and phasic firing patterns, which have been previously reported [46]. These differing patterns are consistent with a heterogeneous population of sensory neurons. The sensitivity of firing to tetrodotoxin (TTX) and lidocaine confirms functional expression of voltage-gated sodium channels, including TTX-sensitive subtypes such as Nav1.7, which are highly expressed in nociceptors. TTX-resistant channels, particularly Nav1.8, which are strongly implicated in neuropathic pain [1; 13; 24], were not directly assessed, although they are known to contribute to tonic firing in DRG neurons [20]. Given the prevalence of tonic firing observed here, the recorded population likely included nociceptive neurons.

### Mechanisms of cannabinoid-mediated inhibition

CBD and THC inhibited neuronal firing without significantly altering resting membrane potential or whole-cell current amplitude, indicating that their primary action is unlikely to involve direct pore block of channels contributing to resting membrane properties. Instead, these findings support modulation of neuronal excitability through CB1 and CB2 receptors and additional non-cannabinoid targets. CBD has been shown to interact with multiple ion channels and receptors, including voltage-gated sodium channels, TRP channels, GIRK channels, and 5-HT receptors [3; 6; 40; 43]. Consistent with this, no direct modulation of Nav1.7 was observed in our assays, suggesting indirect regulation of excitability.

A significant methodological observation was that CBD and THC effects on DRG neurons were irreversible in conventional solvents (DMSO, ethanol), but reversibility was restored by inclusion of bovine serum albumin (BSA), likely due to reduced adsorption of lipophilic compounds to surfaces [30]. This improved experimental reproducibility and enabled repeated recordings.

### Evidence for receptor-specific and non-specific actions

WIN55,212-2, a cannabinoid receptor agonist, did not significantly alter neuronal firing under these conditions. However, co-application of CBD with WIN55,212-2 enhanced the inhibitory effect on neuronal activity compared with CBD alone, suggesting a functional interaction between receptor activation and CBD-mediated mechanisms.

The inhibition of neuronal firing by the CB2-selective inverse agonist JTE-907 suggests a functional role for CB2 receptors in modulating DRG neuronal excitability. Although CB2 receptors are predominantly expressed on immune cells [4], these findings indicate that they may contribute, directly or indirectly, to neuronal regulation in this context.

### Terpene-mediated modulation of neuronal excitability

In addition to cannabinoids, terpene mixtures demonstrated robust inhibition of neuronal firing. Individual terpenes, such as linalool, β-pinene, and myrcene, produced similar inhibitory effects, although their actions were not readily reversible, likely reflecting prolonged membrane retention due to their lipophilic nature. These findings are consistent with previous studies demonstrating analgesic properties of terpenes in animal models [2; 28].

Terpenes likely act through multiple targets, including ion channels and inflammatory signalling pathways [23; 36], although their precise mechanisms remain incompletely defined. In addition to direct neuronal effects, their ability to modulate inflammatory processes may contribute to reduced nociceptive signalling in chronic pain states.

### Synergistic interactions

Combined application of CBD and THC produced more rapid and pronounced inhibition than either compound alone. Individually, both compounds exhibited slower, time-dependent suppression of firing, consistent with their lipophilicity and indirect, G protein–coupled receptor-mediated mechanisms. Co-application accelerated both onset and magnitude of inhibition, suggesting convergence on shared signalling pathways that enhance suppression of neuronal excitability.

A CBD:THC ratio of 1:5 was sufficient to abolish action potential generation, indicating that lower concentrations may achieve maximal effects when combined. This supports the possibility of enhanced efficacy with combined cannabinoid treatments [11]. Such interactions may have clinical relevance, as reducing THC exposure could limit psychoactive effects while preserving analgesic efficacy.

Terpene mixtures also exhibited enhanced inhibitory effects in the presence of low concentrations of CBD, consistent with previous reports of an “entourage effect” [37]. Together, these findings support the concept that multi-component cannabinoid formulations may provide enhanced efficacy over single-compound approaches, although further work is required to define underlying mechanisms and optimise therapeutic combinations.

### Plant-based cannabinoid

The inhibitory effects observed with whole-plant extracts are consistent with the findings obtained using purified cannabinoids and terpene mixtures. *Cannabis sativa* cultivars exhibit substantial phytochemical diversity, with individual plants containing distinct combinations of cannabinoids, terpenes, flavonoids, and other bioactive constituents. As the present study demonstrated enhanced inhibition of neuronal firing following co-application of cannabinoids and terpenes, it is plausible that the activity of plant extracts reflects interactions among multiple constituents rather than the effects of a single compound alone.

These findings provide further support for the concept of an entourage effect, whereby cannabinoids and terpenes act synergistically to modulate neuronal excitability. Consequently, whole-plant extracts may offer advantages over purified cannabinoid preparations by targeting multiple signalling pathways simultaneously.

In particular, extracts enriched in CBD, while containing lower concentrations of THC, may retain inhibitory effects on nociceptive signalling while reducing the risk of THC-associated adverse effects. Defining the optimal phytochemical composition for analgesia will require further mechanistic studies and controlled clinical trials. Nevertheless, the present findings suggest that carefully characterised plant-derived formulations may provide therapeutic benefits beyond those achievable with individual cannabinoids alone.

### Relevance to neuropathic pain

In experimental models of neuropathic pain, changes in cannabinoid receptor expression have been observed. For example, CB2 receptor expression is upregulated [18], while CB1 receptor levels are reported to decrease in some models [29; 42]. A limitation of the present study is that experiments were conducted in DRG neurons derived from healthy mice, which may not reflect the altered excitability and receptor expression observed in chronic pain states. As such, the pharmacological activity of these compounds may differ under pathological conditions. Compounds targeting CB2 receptors may be particularly relevant in neuropathic pain due to increased receptor expression. In addition, anti-inflammatory effects mediated via CB2 receptors on immune cells may further contribute to analgesia.

### Cardiac considerations

Our findings agree with previous reports demonstrating a greater potency of CBD than THC in inhibiting hERG currents [31]. The terpene mixture also reduced hERG currents, suggesting that terpenes can modulate hERG channel activity under these experimental conditions. Notably, the plant extracts inhibited neuronal firing without significantly affecting hERG currents, raising the possibility of a wider therapeutic window compared with individual compounds. Nevertheless, the translational relevance of these findings remains uncertain and will require evaluation in vivo, where exposure levels, metabolism, and pharmacokinetic factors may substantially influence both efficacy and cardiac safety.

## Conclusion

In summary, this study demonstrates that cannabinoids and terpenes inhibit excitability in peripheral sensory neurons through a combination of receptor-mediated and indirect mechanisms. Their synergistic interactions support the concept of an entourage effect [7], suggesting that multi-component formulations can enhance inhibitory effects on neuronal firing. These findings suggest that plant-derived cannabinoid preparations may offer advantages over single-molecule therapies by targeting multiple pathways simultaneously, potentially achieving greater efficacy at lower doses and improving tolerability. This is particularly relevant for chronic pain, where long-term treatment is required. Notably, CBD–terpene combinations may represent a promising non-psychoactive approach to pain management.

## Supporting information

ESI

## Acknowledgements

The work was supported by 113-Botanicals (grant to JS, BWG, HC).

