## Supplementary material for "Endocannabinoid ligands (CBD, Δ9THC, and Terpenes) inhibit excitability of mouse dorsal root ganglion neurons and exhibit synergistic inhibitory effects": ESI

**Endocannabinoid ligands (CBD, Δ^9^THC, and Terpenes) Inhibit Neuronal Action Potential in Mouse Dorsal Root Ganglion Neurons**

Hedaythul Choudhury^1,2,3*^; Mazin Nicola^3^; Barnaby W. Greenland^4^; Daniel Guest^3,4^; John Spencer^1,4*^; Andrew Dilley^5*^

1. Sussex Drug Discovery Centre (SDDC), School of Life Sciences, University of Sussex, Falmer BN1, 9QJ, UK.
2. Neuroscience, Pharmacology and Physiology (NPP), Division of BioSciences, University College London (UCL), Anatomy Building, Gower Street, London WC1E 6BT, UK.
3. 113 Botanicals, 398 Montrose Avenue, Slough, England, SL1 4TJ, UK.
4. Department of Chemistry, School of Life Sciences, University of Sussex, Falmer BN1 9QJ, UK.
5. Department of Clinical Neuroscience, Brighton and Sussex Medical School, University of Sussex, Brighton, BN1 9PX, UK.

**Figure S1. Whole cell currents are sensitive to block by TEA but not by lidocaine or TTX**

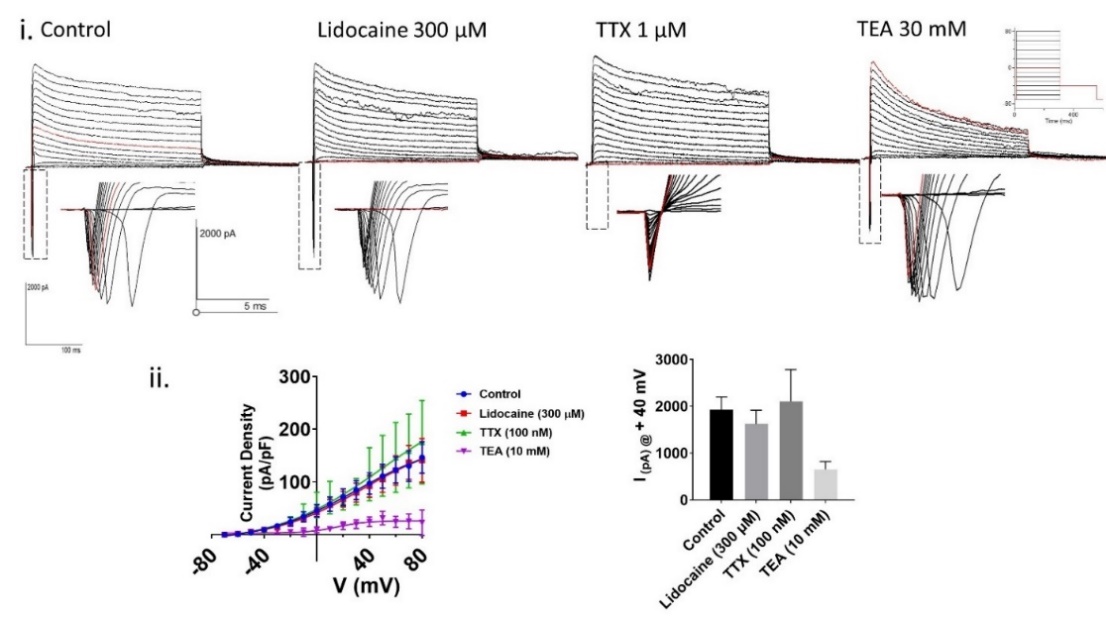

Fig. S1. Example **i)** Effect of Na^+^ and K^+^ channel blockers on whole-cell current. Step currents were activated using the IV step followed by tail protocol (IV protocol: -90 to + 70 mV with 10 mV step followed by tail at -40 mV back to -90 mV, 250 ms duration holding potential -90 mV, 600 ms duration, inter-sweep interval 10s, sampling at 5 Kz. Insert showing the effect of inward current. Application of ion control (n=5), Lidocaine (300 µM) n=4, and TTX (100 nM) n=3, **ii)** Normalised IV currents are plotted against outward current vs voltage on the left graph. The bar graph shows the mean outward current at +40 mV step current.

**Figure S2:** **The effect of BSA and DMSO on mDRG neuronal firing.**

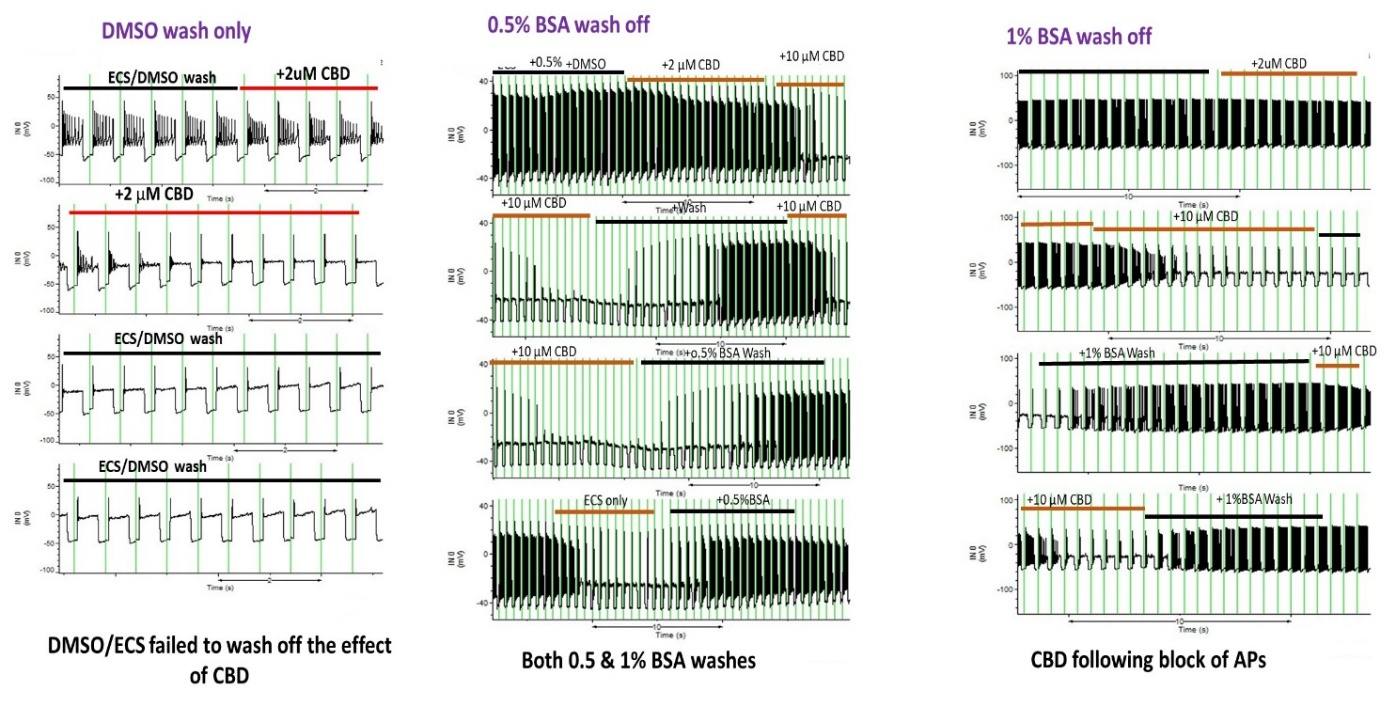

**Figure S2.** Example of whole cell current clamp recording showing the effect of CBD when dissolved in DMSO or BSA on neuronal firing following. Left is the effect of DMSO, middle is 0.5% BSA, and right 1% BSA with ECS solution.

**Figure S3: Effects of CBD and THC on small diameter DRG neurons.**

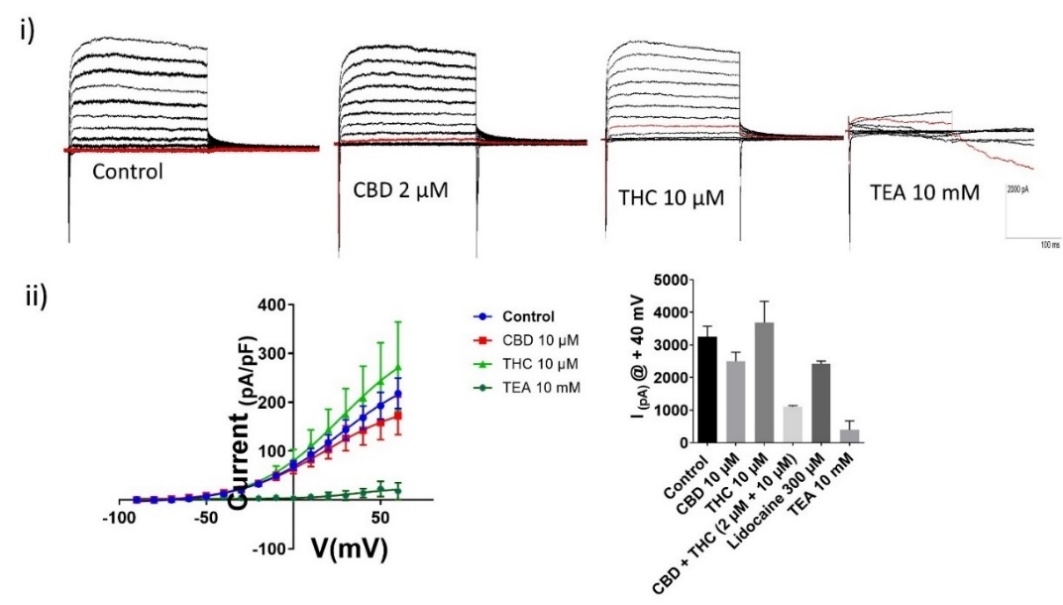

**Figure S3**. Effect of CBD and THC on whole-cell current. **i)** Representative voltage trace recorded from mDRG neurons using IV protocol. **ii)** Normalised IV curves are plotted (current density pA/pF vs voltage mV) using the Boltzmann sigmoidal equation, and V50 (mV) values were calculated**. iii)** Bar graph (right) shows the effect of drugs at +40 mV, and V_50_ was calculated from the curve fit of the Boltzmann sigmoidal equation. Data are presented as Mean ± sem (n).

**Figure S4**.Analysis of HPLC trace showing different cannabinoids in the plant extract

**A.**

**Extract A	Extract B	Extract C**

mAU

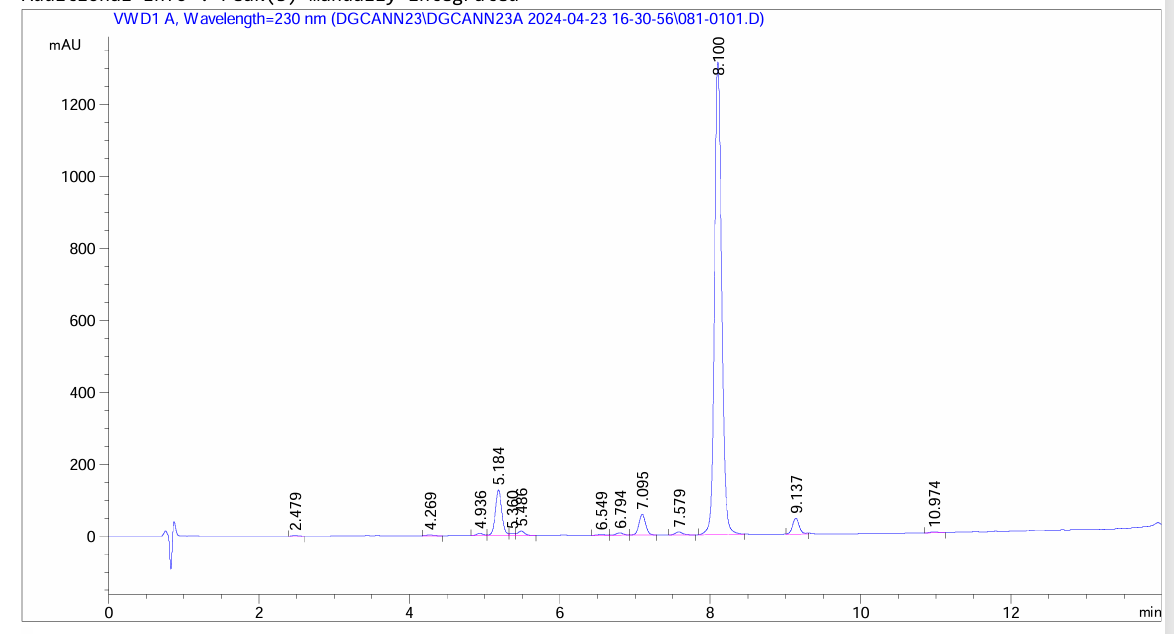

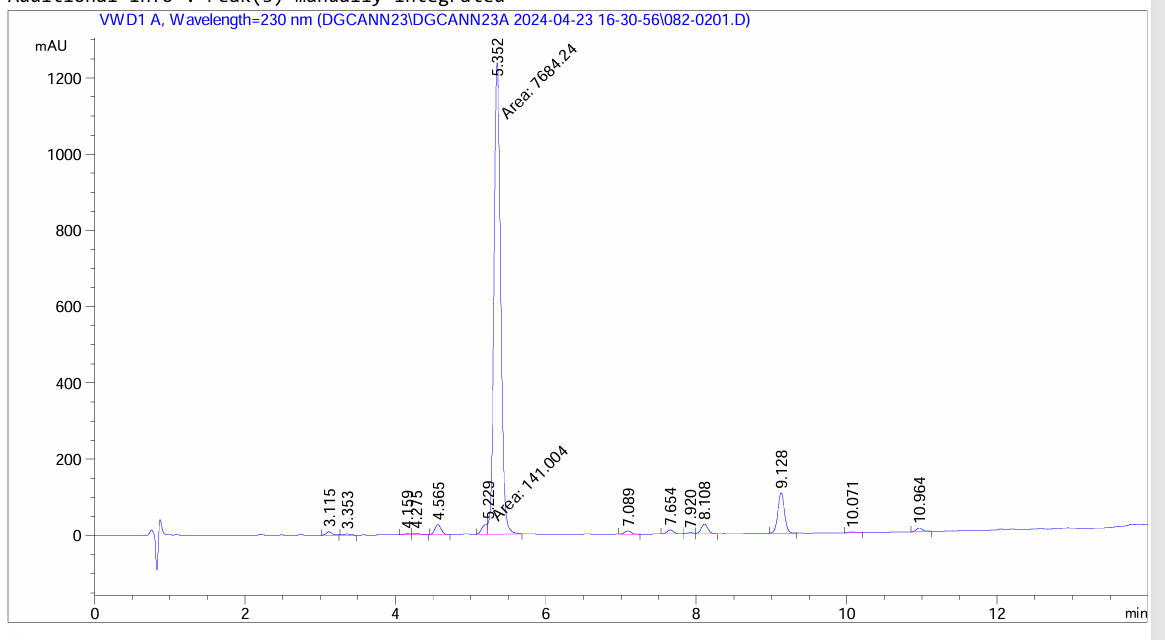

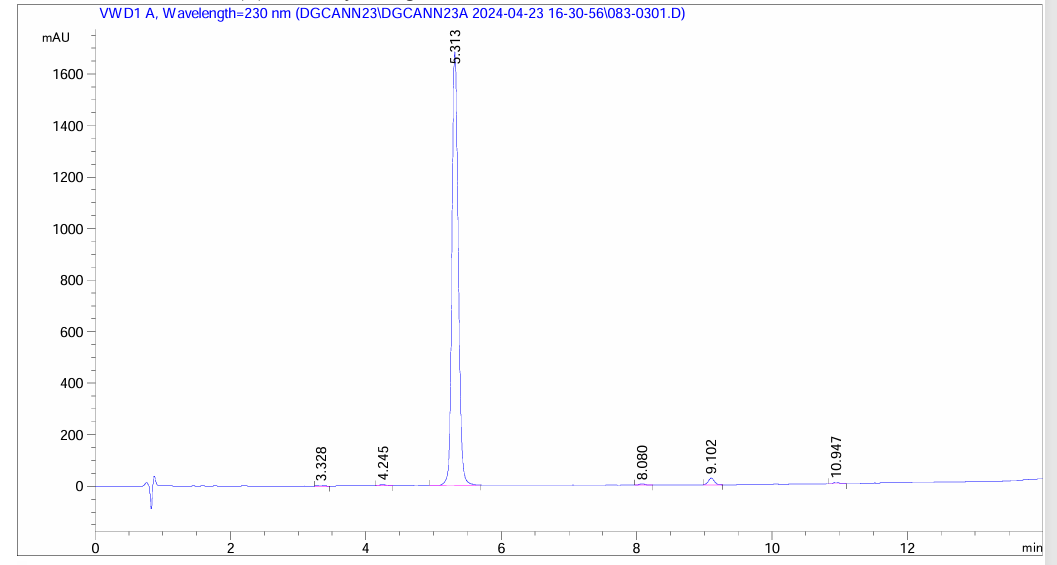

**B.**

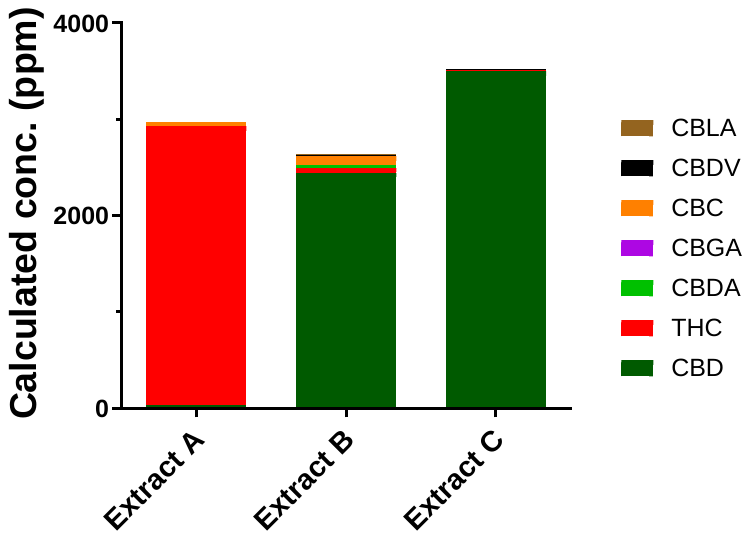

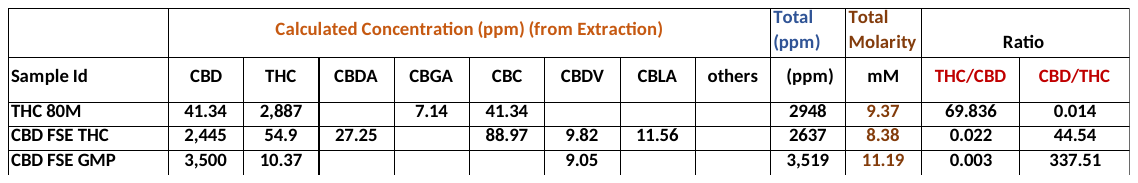

**C.**

Note: THC80M (Extract A), CBD FSE THC (Extract B) and CBD FS GMP (Extract C)

**Figure S4**. A. Showing HPLC and mass spectroscopy data, B. Data table showing different cannabinoids and their concentration either as ppm or ratio of CBD: THC, and finally C. showing bar graph of these three products calculated as ppm.
